# Spectral algal fingerprinting and long sequencing in synthetic algal-microbial communities

**DOI:** 10.1101/2024.07.08.602014

**Authors:** Ayagoz Meirkhanova, Sabina Marks, Nicole Feja, Ivan A. Vorobjev, Natasha S. Barteneva

## Abstract

1. Synthetic biology has made progress in creating artificial microbial and algal communities, but technical and evolutionary complexities still pose significant challenges.
2. Traditional methods for studying microbial and algal communities, such as microscopy and pigment analysis, are limited in throughput and resolution. In contrast, advancements in full-spectrum cytometry enabled high-throughput, multidimensional analysis of single cells based on their size, complexity, and spectral fingerprints, offering more precise and comprehensive analysis than conventional flow cytometry.
3. This study demonstrates the use of full-spectrum cytometry for analyzing synthetic algal-microbial communities, facilitating rapid species identification and enumeration. The workflow involves recording individual spectral signatures from monocultures, utilizing autofluorescence to distinguish them from noise, and subsequent creation of a spectral library for further analysis. The obtained library is used then to analyze mixtures of unicellular cyanobacteria and synthetic phytoplankton communities, revealing differences in spectral signatures. The synthetic consortium experiment monitored algal growth, comparing results from different instruments and highlighting the advantages of the spectral virtual filter system for precise population separation and abundance tracking. This approach demonstrated higher flexibility and accuracy in analyzing multi-component algal-microbial assemblages and tracking temporal changes in community composition.
4. By capturing the complete emission spectrum of each cell, this method enhances the understanding of algal-microbial community dynamics and responses to environmental stressors. With development of standardized spectral libraries, our work demonstrates an improved characterization of algal communities, advancing research in synthetic biology and phytoplankton ecology.

## Introduction

In recent decades, synthetic biology has made significant progress in building artificial microbial and algal communities (Deng et al., 2022; Fu et al., 2020); however, the difficulty of engineering complex biological systems remains significant due to technical hurdles and the intricate nature of biosystems, which evolve and refine themselves over time (Hartwell et al., 1999). Community selection experiments address fundamental questions in ecology and evolution, as well as applied biotechnological issues, yet constructing more complex stable communities to study the effects of different bacteria on community dynamics remains challenging. Some of the challenges include the fact that uni-algal systems are not stable, and bacterial contamination is likely if the algal community is maintained heterotrophically on a medium with a fixed source of carbon (Andersen, 2005) or in the presence of dead cells. Several studies have demonstrated that algal-microbial synthetic consortia, which exhibit mutualism, can be engineered (De-Bashan et al., 2002; Cho et al., 2015; Xu et al., 2016). However, algal-microbial consortia development is usually limited by studying a few algae (often, systems are over-dominated by certain algae) or only the bacterial part of the consortia, with some exceptions (Segev et al., 2016; Kolter, 2024). Moreover, long-term homeostasis of algal-microbial consortia could be difficult to maintain since evolutionary constraints are known to limit the long-term stability of synthetic systems (Castle et al., 2021), and behavior and composition of engineered communities is “unpredictable” (Brenner et al., 2008).

Finally, the absence of methods allowing to follow the dynamics of mixed algal-microbial systems further complicates the development of complex synthetic systems. Identification and characterization of individual microalga and bacteria in the consortium are essential for engineering such consortia and development of subsequent applications (Perera et al., 2019). While microbial community composition analysis has advanced significantly, including classification of reads up to the species level (Curry et al., 2022), effective methods for plankton identification and characterization are still limited. Traditionally, microbiome analysis in such systems was conducted at the genus and class levels due to limitations in next-generation sequencing. However, with the development of long sequencing techniques, more detailed analyses have become possible. For instance, in a long-term mesocosm experiment, species-level visualization using imaging cytometry combined with long sequencing was employed (Meirkhanova et al., 2024). The integration of a full-spectrum approach would, therefore, provide a distinct advantage by enabling more precise and comprehensive analysis.

An algal community can be categorized into subgroups based on morphology, size, cellular functions, and interactive relationships. However, morphology-based taxonomy has limitations, particularly in distinguishing cell types with similar morphologies. Traditional methods for studying microbial and algal communities, such as microscopy and pigment analysis, are constrained by throughput, taxonomic resolution, and functional characterization, primarily differentiating specific taxonomic groups. Spectrofluorimetry, spectroscopy, and HPLC-based approaches provide averaged integrated data at a population level (Poryvkina et al., 1994; Falkowsky, Oliver, 2007; Roy et al., 2011; Swan et al., 2016; Liu et al., 2021). The maximum of taxonomic groups identified by HPLC-pigments analysis typically limited by 4-7 distinct groups (Catlett, Siegel, 2018; Kraemer, Siegel, 2019). Though full-resolution reflectance spectra are used in discriminant and classification analyses of algae by airborne hyperspectral sensors, applying spectral unmixing algorithms based on reference data (Hochberg, Atkinson, 2003; Rodriguez et al., 2017; Kramer, Siegel, 2019; Cause-Nicholson et al., 2021), there is still a need for adequate *in situ* method. Conventional flow cytometry struggles to distinguish cells from other particulate matter in the water matrix, and imaging flow cytometry can differentiate only a few fluorescent channels. In contrast, full-spectrum cytometry has recently transformed the field by offering high-throughput, multidimensional analysis of single cells based on their size, complexity, and spectral fingerprint (Futamura et al., 2015; Robinson, 2022; Dott et al., 2024).

This study introduces full-spectrum cytometry as a powerful tool for analyzing synthetic algal-microbial communities, facilitating rapid, high-throughput analysis from species identification to enumeration without extensive sample preparation. By capturing the complete emission spectrum of each cell, full-spectrum cytometry provides valuable insights into phytoplankton community composition, structure, and dynamics, as well as their responses to environmental stressors that influence community structure (Barteneva et al., 2019, 2023; Jeppesen et al., 2005). Algal cells, distinguished by autofluorescent spectral signatures, are separated taxonomically, while light scattering parameters assess cell size and complexity, distinguishing target events from debris. Spectral cytometry identifies highly autofluorescent subpopulations using virtual filtering (Barteneva et al., 2019) or only autofluorescence finder algorithm (Wanner et al., 2022). Characterizing microbiota associated with algae from in-house cultures or natural consortia is crucial as microalgae growth phases correlate with shifts in microbial phylotypes (Geng et al., 2016), although microbial dynamics in synthetic systems remain understudied. This research demonstrates that modern spectral cytometry, coupled with species-level 16S rRNA sequencing, effectively identifies and enumerates algae based on their spectral signatures, advancing our understanding of phytoplankton ecology, synthetic biology, and evolution.

## Methods

### Sample preparation and instrument setup

Algal cultures (**Suppl. Table 1**) were obtained from Culture Collection of Algae at the University of Göttingen (Germany) and Central Collection of Algal Cultures at the University of Duisburg-Essen (Germany). Cultures were maintained in Basal Medium (BM) at 21°C, under 12/12 L/D cycle in algae growth chamber AL-30L2 (Percival, USA). Samples were recorded in parallel on three instruments: 7-laser ID7000 Spectral Cell Analyzer (Sony Biotechnology, USA), 5 laser BD FACSAria Special Order cell sorter (BD Biosciences, USA) and 4-lasers ImageStreamX MarkII (Amnis - Cytek, USA) imaging flow cytometer equipped with 40X and 60X objectives. Samples prepared for cytometric analysis included mono- and mixed cultures and were recorded using a standard 24-tube rack. Each sample was placed in round-bottom 5mL tubes, with a minimum volume of 250µL, and arranged in specified positions on the rack. Prior to data acquisition, instrument settings were adjusted and set in accordance with each sample (FSC and SSC gains depending on algal culture, threshold value – 11%, PMT voltage ranging from 40% to 70%). Samples were recorded at minimum flow rate (=1) and 50,000 events were set for data acquisition. FSC_A vs SSC_A plots were used for the initial assessment of instrument settings in the “Preview” mode prior to recording.

Next, synthetic consortium was set up and included representatives of three major phytoplankton groups – Cyanobacteria (*Microcystis* sp., *Synechococcus* sp., *Synechocystis* sp.), Chlorophyta (*Chlamydomonas* sp., *Scherffelia* sp., *Haematococcus pluvialis*) and Cryptophyta (*Rhodomonas* sp., *Cryptomonas* sp., *Chroomonas* sp.) – with size ranging from 2 to 25 µm (**Table 1, Suppl. Fig. 1**), growth of which was monitored daily for the period of 9 days. Triplicates were also analyzed in parallel on BD FACSAria cell sorter (BD Biosciences, USA) and ImageStreamX MarkII (Amnis - Cytek, USA) imaging flow cytometer. Microbial community composition during the last day of the experiment was assessed using nanopore-based full-length 16S rRNA gene sequencing.

**Table 1.**
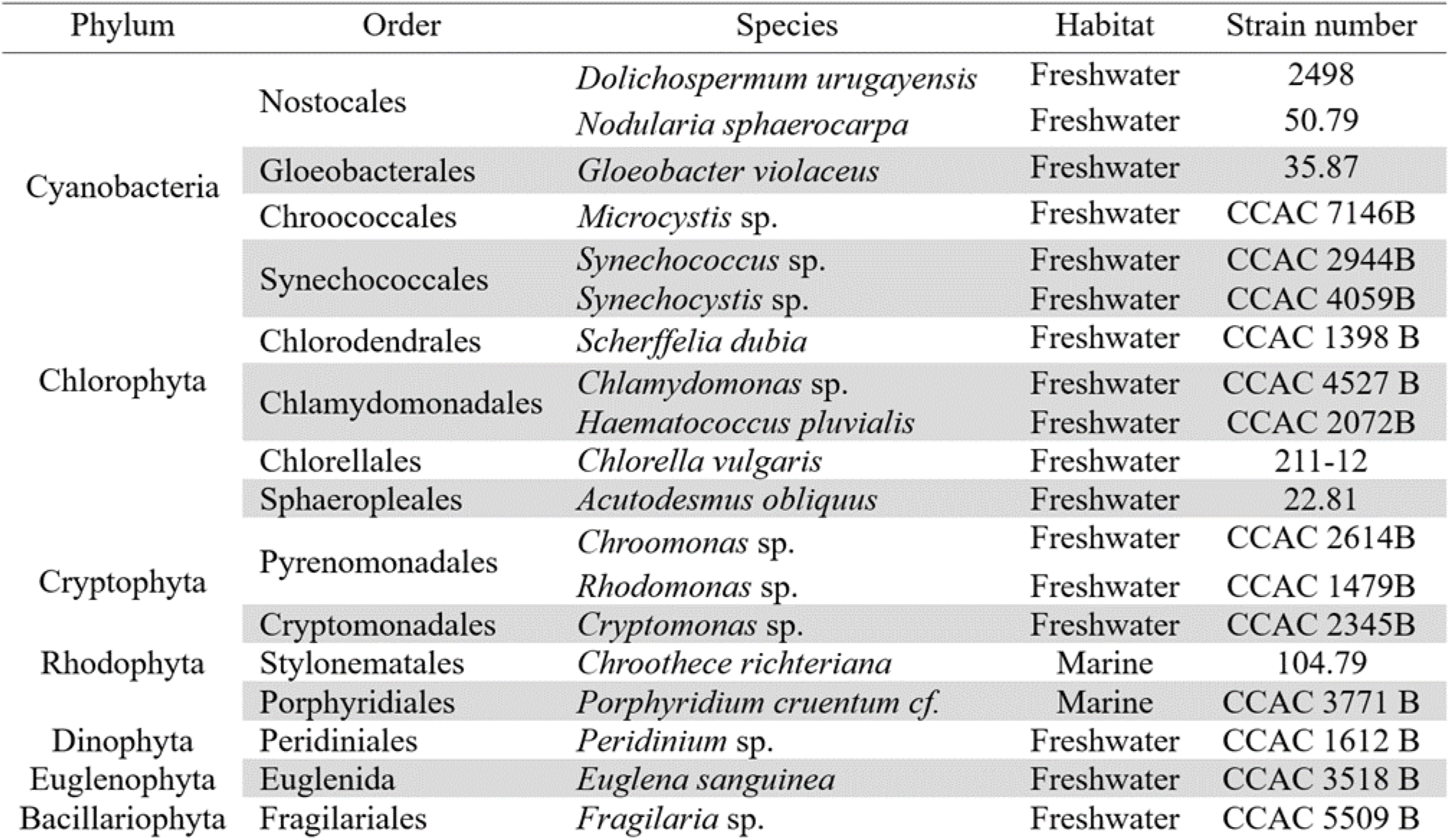
List of algal cultures used for recordings during artificial mix experiment and library construction using ID7000 spectral cell analyzer.

### Autofluorescence Finder Tool and Virtual Filters setup

The ID7000 software’s Autofluorescence Finder Tool was used to analyze the algae samples, treating autofluorescence as an independent fluorescent parameter. This feature is crucial for the taxonomic identification of different algae. By incorporating excitation lasers ranging from 320nm to 808nm, we included unique emission regions in the algal spectral signature (fluorescence intensity and shape), distinct from the chlorophyll peak and highly variable among different algae. This setup allowed for the detection of individual autofluorescent spectral signatures for each of the recorded algal monocultures. Following the identification of the best separation between autofluorescent populations, subpopulations of interest were gated and examined for their spectral signatures. Each population demonstrated a unique autofluorescence spectrum. Furthermore, Autofluorescence Finder tool was used for virtual filters setup. In a similar manner, highly variable regions between different algal taxonomic groups are defined and used to set a pair of virtual filters (VFs). Defined VFs are next used to separate subpopulations of interest within multi-component mixes.

### Data analysis

Initial cytometric data analysis and population identification were performed using the ID7000 software. Identification of subpopulations within an artificial mix involved consecutive gating steps using a combination of 2D plots with specified virtual filters for best separation. In addition, t-Distributed Stochastic Neighbor Embedding, otherwise known as t-SNE, was done using FCS Express (De Novo Software, USA). A modified version of t-SNE - opt-SNE, a tool allowing the mapping of high-dimensional cytometry data onto a two-dimensional plot while conserving the original high-dimensional structure, was used to independently identify subpopulations within the artificial mix, minimizing gating bias. Obtained clusters were then used to compare data acquired on different cytometers. To display fluorescence intensity dispersion, coefficient of variation was calculated and plotted across all detection channels.

### Sequencing library preparation and bioinformatic analysis

Library preparation, the sequencing run on the MinION Mk1C device and bioinformatic analysis were performed in laboratory following the end of 9-day synthetic consortium experiment. Firstly, DNA was extracted using the DNEasy Power Water Kit (Qiagen, USA) according to the manufacturer’s instructions, with an additional heating lysis step. The full-length 16S gene was then amplified using the universal 16S primer pair 27F (5′-AGAGTTTGATCCTGGCTCAG-3′) and 1492R (5′-TACGGYTACCTTGTTACGACTT-3′). Library preparation followed the manufacturer’s protocol (ONT, UK). The PCR reaction mixture included nuclease-free water, input DNA, LongAmp Hot Start Taq 2X Master Mix (NEB, USA), and the respective 16S barcode primer. PCR products were next purified using AMPure XP beads (Beckman Coulter, USA). In the final library preparation step, sequencing adapters were added to the barcoded samples. Raw data underwent basecalling using the guppy basecaller (ONT, UK). The resulting FASTQ files were processed using Python algorithms beginning with read assessment, followed by quality (Q-score>10) and read length filtering. The next steps involved adapter sequence removal and demultiplexing of reads into their respective barcodes. The demultiplexed reads were then classified using the Emu taxonomic abundance estimator (Curry et al., 2022) along with a custom reference database (Len et al., 2023) up to the species level (https://gitlab.com/treangenlab/emu). Lastly, the data were rarefied prior to further analysis.

## Results

The first step in the analysis of complex communities requires construction of a library with unique spectral signatures for different phytoplankton groups. For this purpose, single phytoplankton cultures were recorded first on ID7000 Spectral cell analyzer (Sony Biotechnology), and individual spectral signatures were then extracted. **Fig. 1** demonstrates the general workflow for the analysis of *Microcystis* sp. monoculture as an example. During the recording, emission spectra across 7 lasers were obtained, from algal cultures with presence of background noise/cellular debris (**Fig. 1a**).

**Figure 1.**
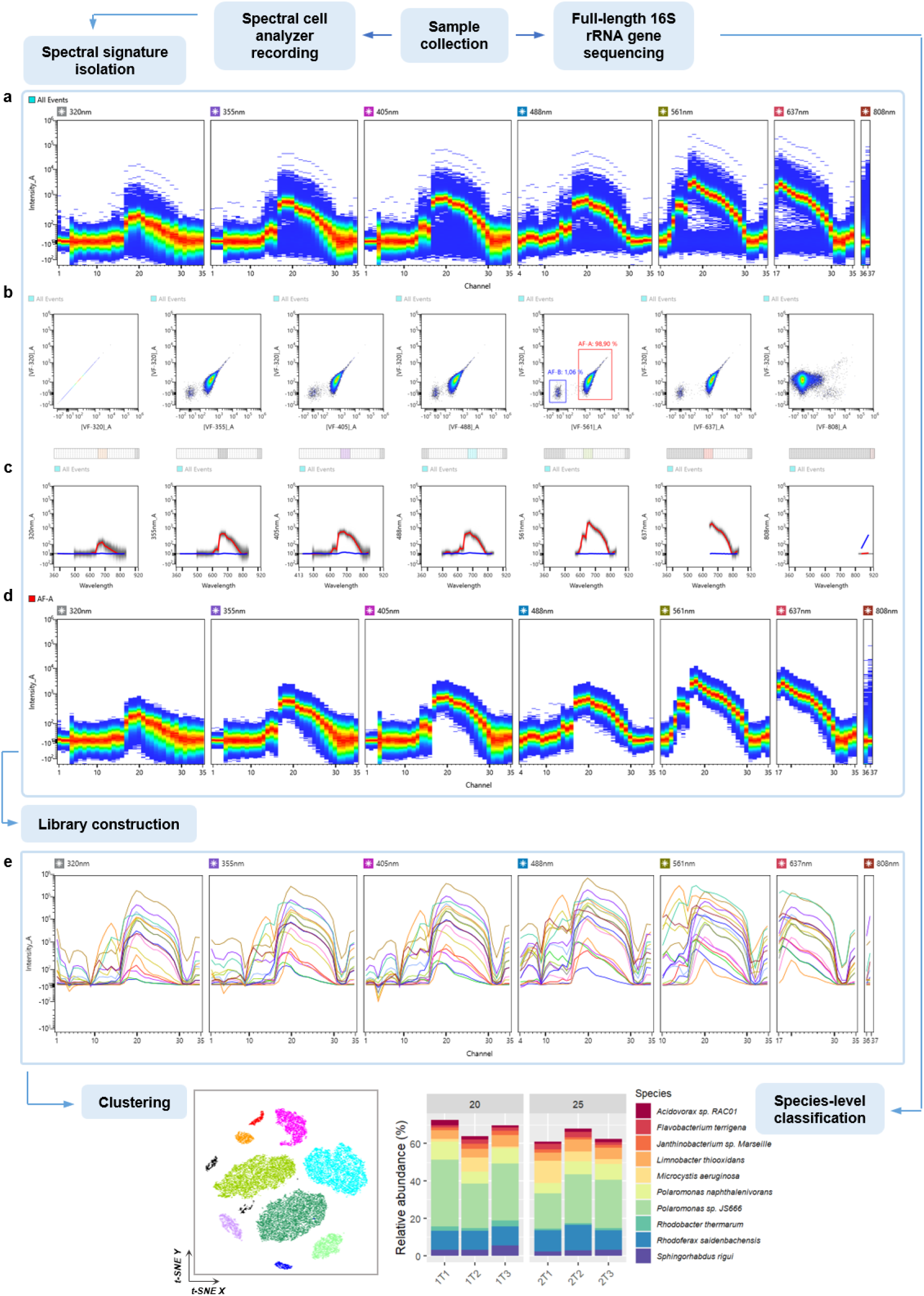
Schematic overview of the proposed experimental workflow for the analysis combining full-spectrum cytometry and sequencing. The workflow involves parallel analysis of obtained samples using MinION Mk1C-based sequencing platform (ONT) and ID7000 Spectral cell analyzer (Sony Biotechnology). The first step in the analysis of spectral data requires **(a)** recording and isolation of single spectral signatures, for which “Autofluorescence Finder” tool is utilized. During this step **(b)** a set of optimal virtual filters for discrimination of autofluorescent populations is determined. As determined using the tool, **AF-A** corresponds to *Microcystis* sp. and **AF-B** to debris within the sample; **(c)** respective emission spectra of two defined autofluorescent populations displayed below (red line – **AF-A**, blue line – **AF-B**). **(d)** Single spectral signature is then extracted and **(e)** a library, containing unique spectral signatures corresponding to respective phytoplankton species, is then constructed and utilized during subsequent analysis, including unsupervised clustering techniques (e.g., t-SNE).

Autofluorescence was then utilized in order to discriminate the spectra of interest from the debri/noise. In some cases, direct gating based on light scattering is possible as well, however, for picoplankton (e.g., *Synechococcus*), with cell size ranging from 0.2 to 2 µm, the gating is not precise. Instead, intrinsic cell fluorescence was used to distinguish events of interest from low-intensity/no-intensity events. With “Autofluorescence Finder” tool clear separation between the two autofluorescent populations was observed across all lasers, except for 808 nm, given the optimal set of virtual filters (VFs). **Fig. 1b,c** demonstrates pairwise density plots of [VF-320]_A against VFs of 6 remaining excitation lasers as an example, with detection of two populations: AF-A – *Microcystis* sp. – 98.9% and AF-B – debris – 1.06% ([VF-320]_A vs. [VF-561]_A). In this case, filters were set around channel 20, where emission intensity was the highest. Unique spectral signature for *Microcystis* sp. was then extracted by plotting the emission of autofluorescent population AF-A on a ribbon plot (**Fig. 1d**). The same workflow was applied for the acquisition of spectral signatures of the remaining algal monocultures (**Figure 1e**), resulting in a spectral library used for further analysis.

To discriminate multiple components in a suspension, a mixture of unicellular cyanobacteria was used and recorded for further analysis (**Fig. 2**). In addition, components of the mix represented variable taxonomic groups at the Order level (*Microcystis* sp., belonging to Chroococcales), as well as members of the same Order (*Synechococcus* sp., *Synechocystis* sp., belonging to Synechococcales). Individual spectral signatures were recorded prior to the mix; **Fig. 2b** demonstrates overlay plot of the acquired spectra across 7 excitation lasers. As seen from the plot, *Microcystis* sp. (red) and *Synechococcus* sp. (blue) possess nearly identical spectral signatures, with slight differences in fluorescence intensity, while distinct region around channels 10-16 (566-646 nm) in the spectra for *Synechocystis* sp. (violet) was recorded. Based on the obtained information, set of optimal VFs was chosen to firstly discriminate autofluorescent populations from unwanted events, and to further separate these populations into respective picoplanktonic groups. Gating strategy is presented in **Fig 2c**, where a combination of [VF-647]_A vs [VF-488]_A reveals two subpopulations, corresponding to: *Synechocystis* sp. and population “A”. Due to the similarity in the spectra of *Microcystis* sp. and *Synechococcus* sp. subpopulation “A” was then differentiated based on SSC values. Identified populations could not be discriminated based on FSC and SSC only (**Fig. 2d**). At this point, the maximum number of individual phytoplankton species that could be discriminated reached 14 (**Fig. 3**).

**Figure 2.**
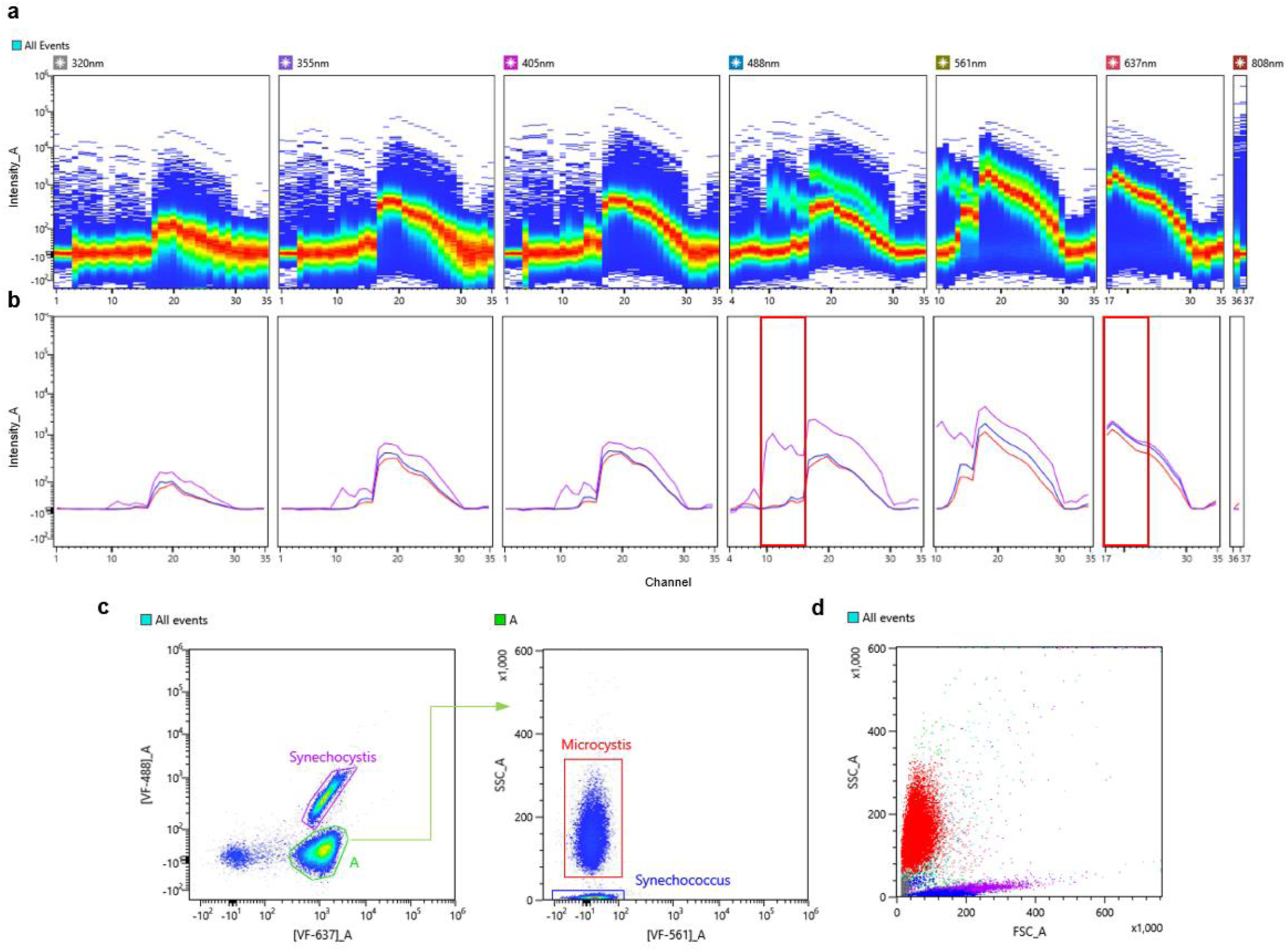
Discrimination of cyanobacteria in a synthetic mix based on autofluorescence. **(a)** Ribbon plot demonstrating combined spectral signatures of three cyanobacterial representatives across 7 excitation lasers: *Synechococcus* sp., *Synechocystis* sp., and *Microcystis* sp. **(b)** Overlay plot of individual spectral signatures for each cyanobacterial representative (*Synechococcus* sp. - blue, *Synechocystis* sp. - violet, and *Microcystis* sp. - red); red regions indicate regions of interest, used for setting an optimal pair of VFs. **(c)** Dot plot of **[VF-637]**_A against **[VF-488]**_A with two autofluorescent populations identified: *Synechocystis* sp., population **A** was then plotted with **[VF-561]**_A against **SSC**_A to resolve two subpopulations *of Microcystis* sp. and *Synechococcus* sp. **(d)** Dot plot demonstrating relative position of each of the identified cyanobacterial species on FSC_A against SSC_A plot.

**Figure 3.**
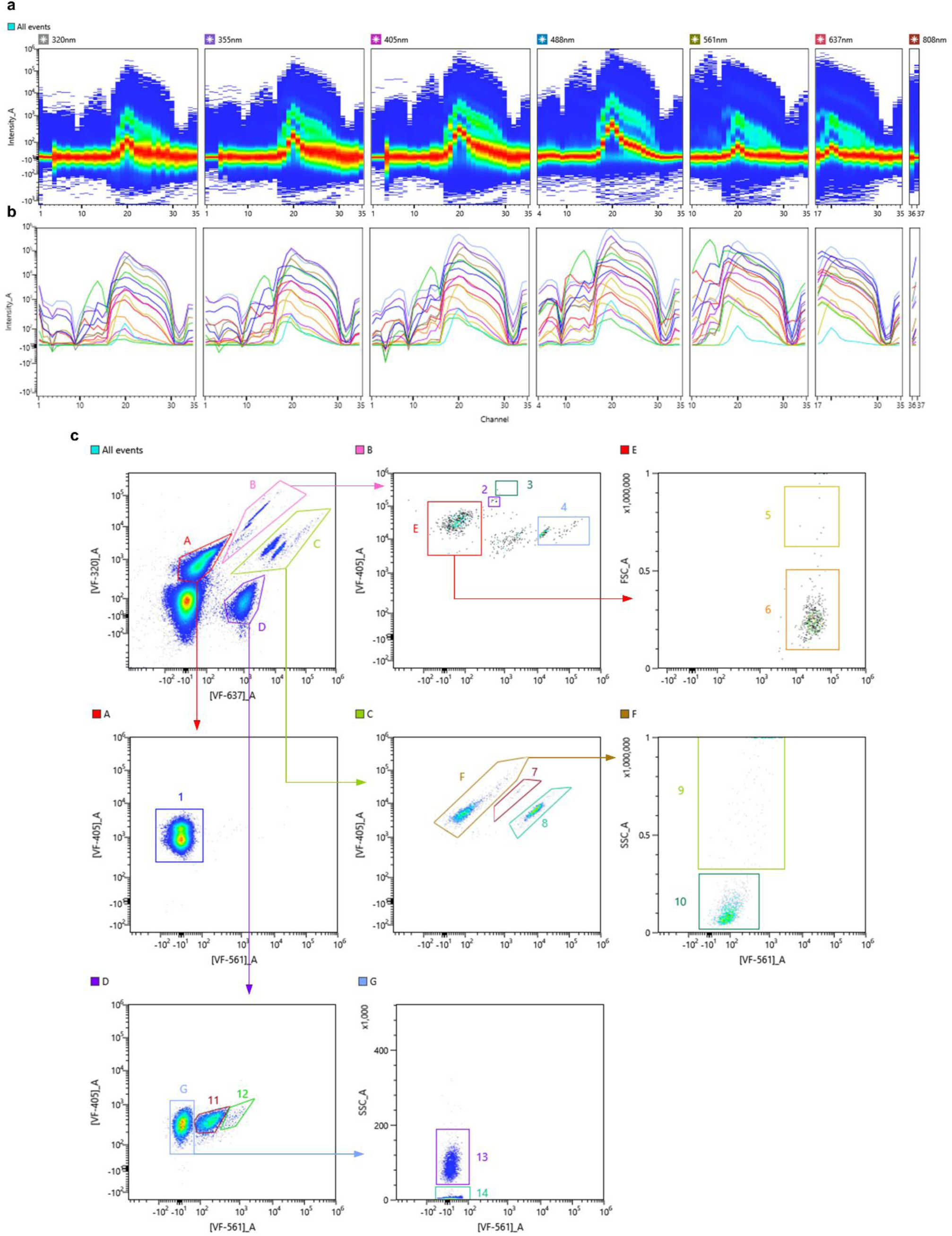
Gating strategy for discrimination of 14 groups of phytoplankton species based on autofluorescence in a complex synthetic mix. **(a)** Ribbon plot demonstrating combined spectral signatures of 16 phytoplankton species across 7 excitation lasers. **(b)** Overlay plot of individual spectral signatures for each phytoplankton representative. (c) Gating strategy for discrimination of each subpopulation within the synthetic mix: 1 – Chlorophyta (*Chlorella vulgaris* and *Acutodesmus obliquus*), 2 – *Euglena sanguinea*, 3 – *Peridinium* sp., 4 – *Cryptomonas* sp., 5 – *Fragilaria* sp., 6 – *Haematococcus pluvialis*, 7 – *Chroothece richteriana*, 8 – *Porphyridium cruentum* cf., 9 – filamentous Cyanobacteria (*Dolichospermum urugayensis* and *Nodularia sphaerocarpa*), 10 – *Chroomonas* sp., 11 – *Synechocystis* sp., 12 – *Gloeobacter violaceus*, 13 – *Microcystis* sp., 14 – *Synechococcus* sp.

Next, applicability of full-spectrum cytometry in monitoring of dynamics of synthetic phytoplankton consortium was assessed. For this purpose, an artificial mix experiment was set up; synthetic mix included 3 representatives of major phytoplankton groups – Cyanobacteria, Chlorophyta and Cryptophyta. Firstly, a library, containing individual spectral signatures covering each phytoplankton species within the mix was constructed. **Fig. 4a** represents 9 spectral signatures across three phytoplankton groups: Cyanobacteria (*Microcystis* sp., *Synechococcus* sp., *Synechocystis* sp.), Chlorophyta (*Chlamydomonas* sp., *Scherffelia* sp., *Haematococcus pluvialis*) and Cryptophyta (*Rhodomonas* sp., *Cryptomonas* sp., *Chroomonas* sp.). As seen from the graph, regions unique for each of the group are present, specifically between channels 10-15 (566-646nm) of the spectra. This information was further utilized to define a set of optimal VF pairs to discriminate these populations in a complex mix. VFs were defined in a way to cluster subpopulations within the same group, while maintaining the difference between the groups. Set of filters [VF-320]_A vs [VF-561]_A (**Fig. 4b**) was chosen as an optimal pair for the first step within the gating strategy (**Fig. 4c**), where three groups are identified and discriminated from the rest of the unwanted events (background noise/cellular debris). In addition to cytometric analysis, samples collected at the end of the experiment were also subjected to sequencing-based analysis.

**Figure 4.**
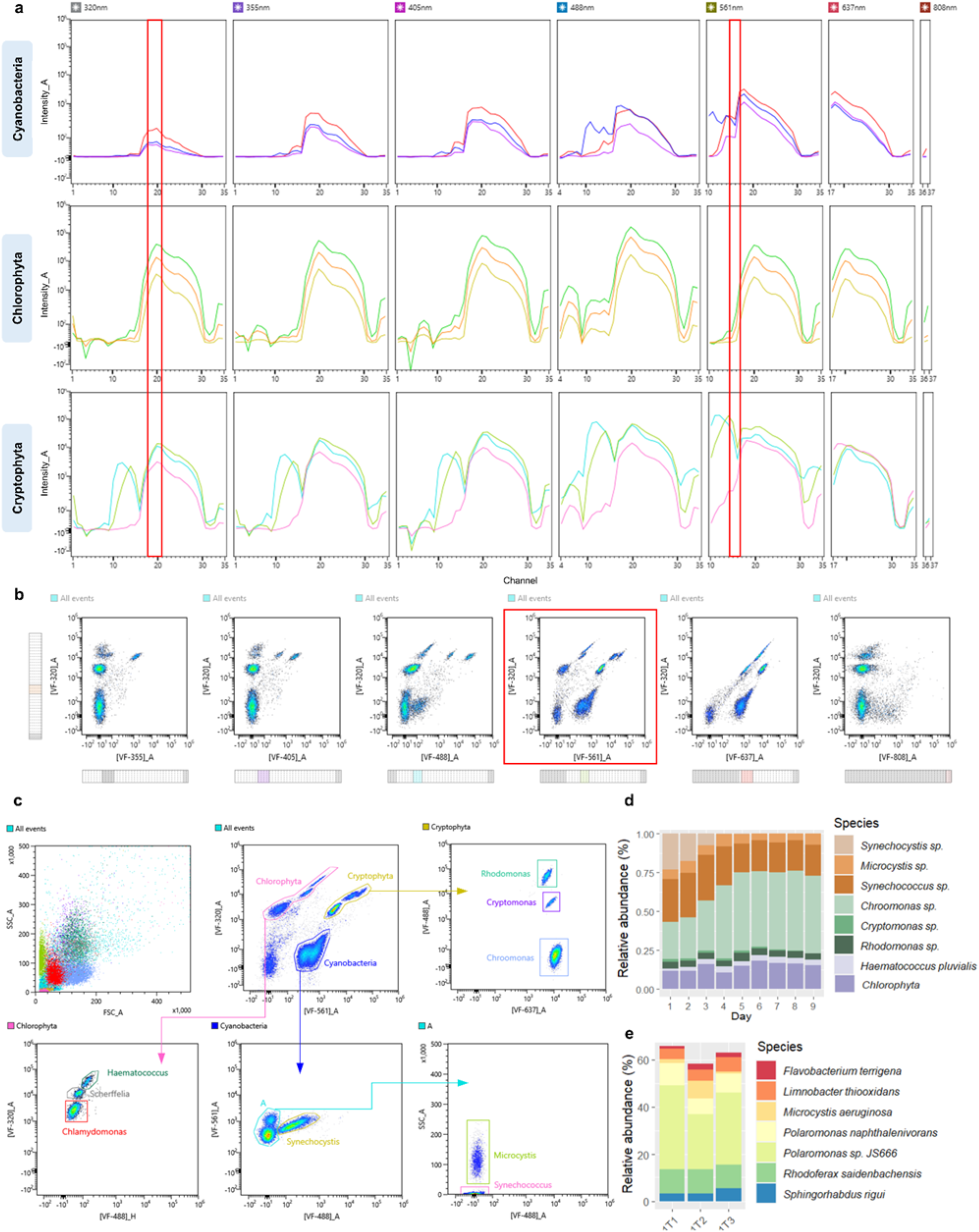
Tracking temporal changes in synthetic community composition. **(a)** Overlay plot of individual spectral signatures for each phytoplankton group representative (Cyanobacteria: violet – *Microcystis* sp., red – *Synechococcus* sp., blue – *Synechocystis* sp.; Chlorophyta: yellow – *Scherffelia dubia*, orange – *Chlamydomonas* sp., green – *Haematococcus pluvialis*; Cryptophyta: pink – *Chroomonas* sp., blue – *Rhodomonas* sp., green – *Cryptomonas* sp.); red regions indicate regions of interest, used for setting an optimal pair of VFs. (**b)** “Autofluorescence Finder” tool with VFs set to regions of interest, demonstrating the best separation of subpopulations on [VF-320_A] vs [VF-561_A] plot. **(c)** Gating strategy, composed of sequential sub-gating steps, used to identify phytoplankton subpopulations within the synthetic community. **(d)** Changes in relative abundance of the artificial mix subpopulations during the nine-day experiment evaluated with ID7000 spectral cytometer. **(e)** Microbial community composition during the last day of the experiment in triplicates at species level revealed through full-length 16S rRNA sequencing.

**Fig. 4d** summarizes averaged changes in the relative abundance of the components of the artificial mix throughout the experiment. Cryptophytes, specifically *Chroomonas* sp., were recorded to be the dominant group within the mix starting from the fourth day, outnumbering cyanobacterial representatives. In addition to cytometric analysis, samples collected at the end of the experiment were also subjected to sequencing-based analysis. **Fig. 4e** demonstrates microbial community composition across three replicates at species levels. Three most abundant phyla besides Cyanobacteria were Proteobacteria and Bacteroidetes, with *Polaromonas sp. JS666* being the most abundant bacterial species. The results showed stability of bacterial composition in artificial algal-microbial mix during 9-days experiment resulting in unsignificant differences between sequenced triplicates.

The next steps involved pairwise combinations of defined filters to discriminate the components of each of the subpopulations. As mentioned previously, due to the high similarity of spectra between *Microcystis* sp. and *Synechococcus* sp., SSC was used to differentiate these populations. A comparison of the data obtained on three instruments revealed limitations of some of the instruments (**Fig. 5**). Opt-SNE dimensionality reduction algorithm was utilized to minimize “gating bias” and group data into clusters with similar characteristics in low-dimensional space. Three datasets were compared: synthetic mix data obtained from ID7000 instrument using a set of pre-defined VFs and a set of VFs matching the filters used on FACSAria, and lastly, data obtained from FACSAria. Both sets of filters utilized on the ID7000 resulted in clear separation of all 9 subpopulations within the mix, while a single population of *Scherffelia* sp. was not possible to discriminate from the rest on the FACSAria instrument. These results demonstrate the higher flexibility a virtual filter system offers in the analysis of multi-component mixes. In addition to more precise population separation obtained with the ID7000 instrument, it was possible to track the abundance of nearly all mix components, as demonstrated in **Suppl. Fig. 2**. Overlay of temporal, spectral signatures recorded for the period of 9 days for each mix member was used to assess the extent of signal intensity dispersion. As revealed through the coefficient of variation, regions most susceptible to differences in intensity generally included channels 1-13 (413.6-612.6 nm) and 30-34 (785.4-835.1 nm). High variation within these regions was also observed when comparing single spectral signatures between excitation lasers (**Fig. 6**).

**Figure 5.**
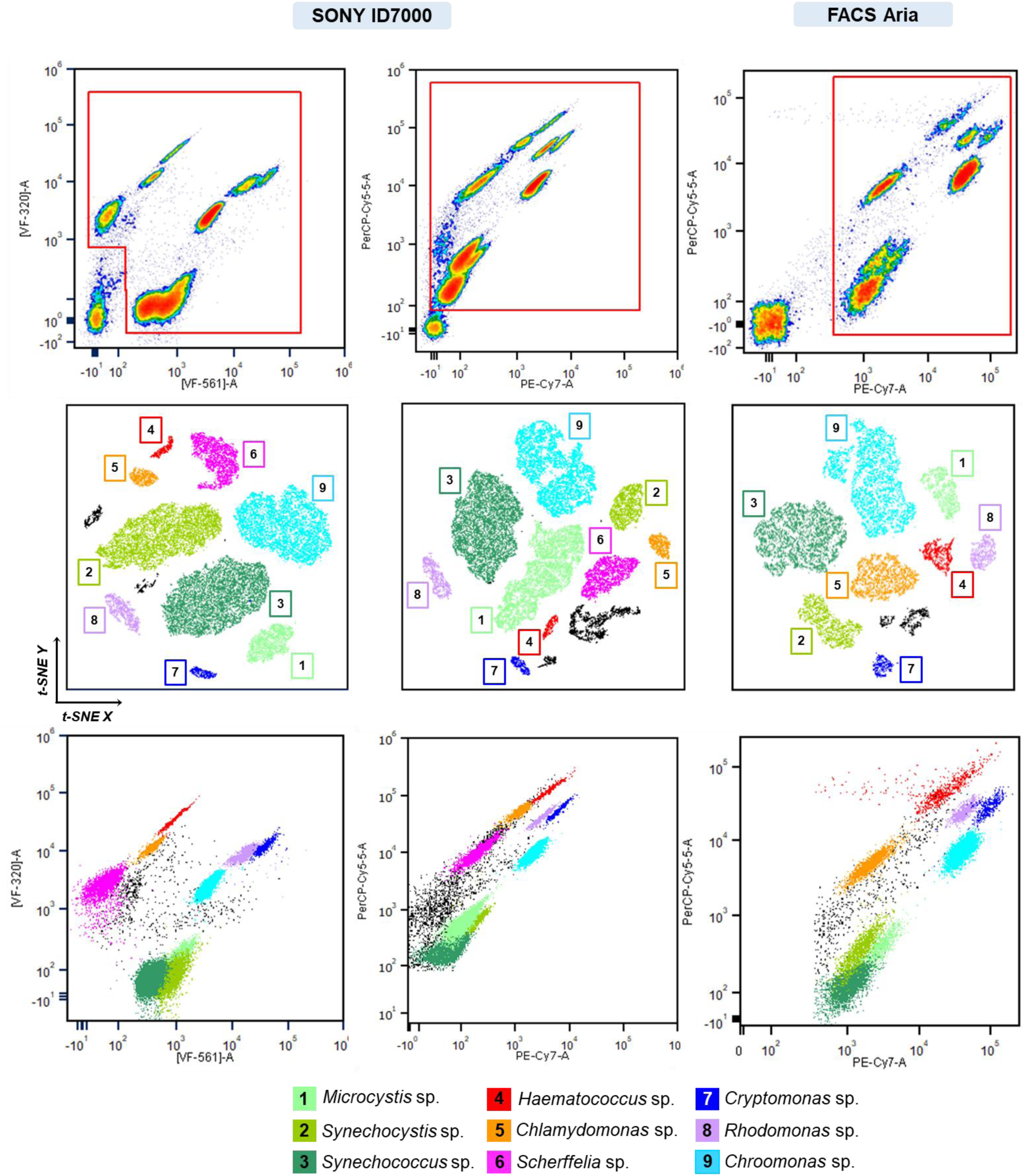
Identification of artificial mix subpopulations compared between SONY ID700 Spectral cell analyzer and BD FACSAria II cytometer. First column represents data recorded on SONY ID7000 using a set of VFs best optimized for artificial mix discrimination. Opt-SNE demonstrates successful discrimination of the 9 subpopulations within the mix based on VF data. Second column represents data recorded on SONY ID7000 using a set of VFs matching optical filters used on BD FACSAria. Similarly, all 9 subpopulations were discriminated. Last column represents data recorded on BD FACSAria. Based on optical configuration of the instrument, only 8 subpopulations were identified using opt-SNE.

**Figure 6.**
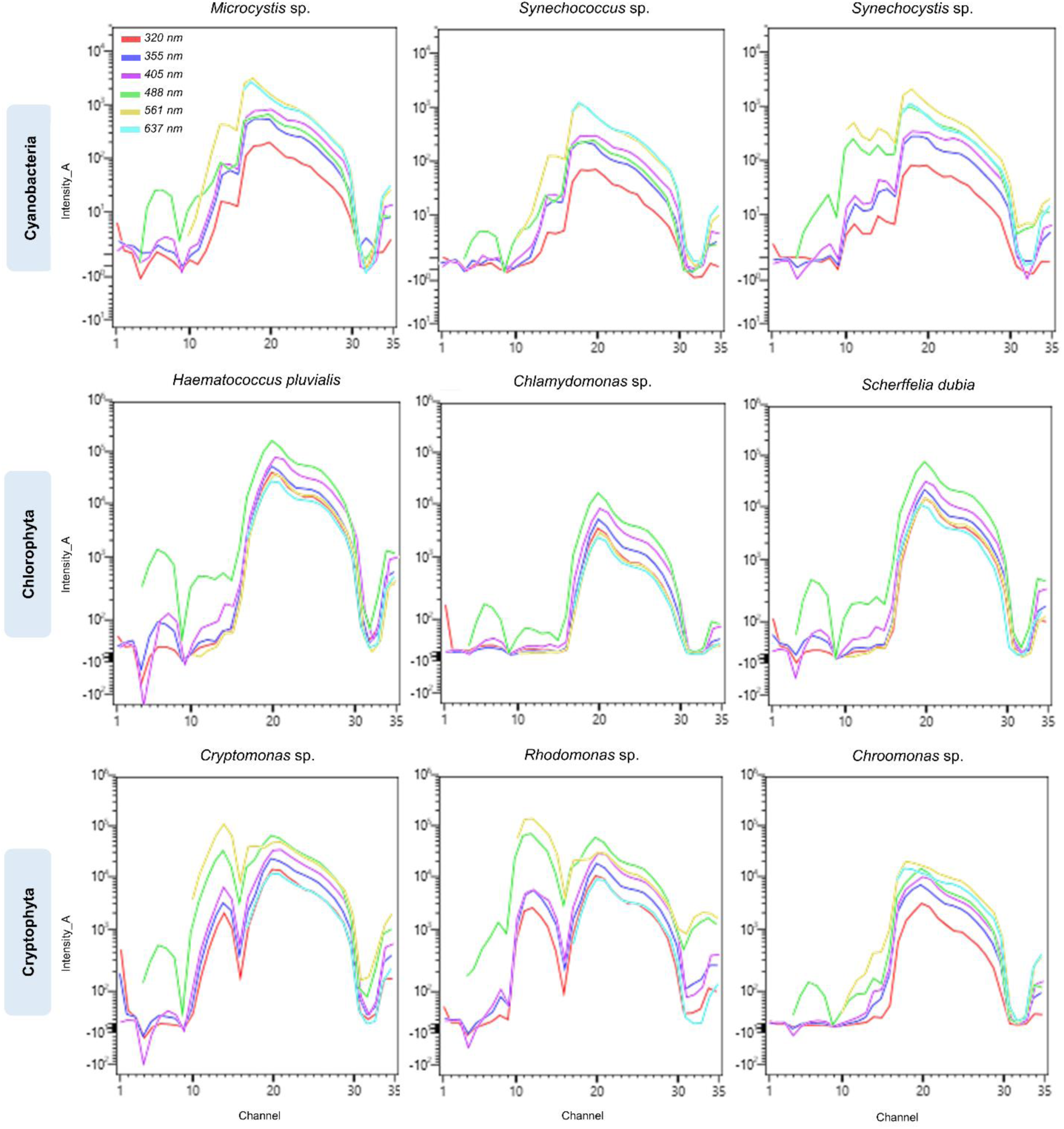
Overlay plots for each phytoplankton species within synthetic mix across 7 excitation lasers to demonstrate variability of emission signal for each excitation source. The first row demonstrates overlay plots for Cyanobacterial representatives, second – Chlorophyta and third – Cryptophyta.

## Discussion

Phytoplankton are extraordinarily diverse, comprising various phylogenetic groups such as diatoms, dinoflagellates, haptophytes, and cyanobacteria. Developing spectral-based technologies for rapidly evaluating the dynamics of microalgal communities is crucial for ecological monitoring and engineering artificial microbial and algal communities. Flow cytometry, a well-established tool in immunological and oncological research, has also been applied to phytoplankton analysis (Trask et al., 1982; Yentsch et al., 1983; Sosik et al., 2010). However, conventional flow cytometers, which rely on narrow band-pass filters to detect specific fluorophores, are limited by spectral overlap and the number of distinguishable fluorophores. Advancements in cytometry, particularly full-spectrum cytometry, present significant improvements in studying synthetic algal-microbial communities. This approach addresses limitations in traditional methods such as microscopy, pigment analysis, and conventional flow cytometry by enabling rapid, high-throughput, and label-free analysis of single algal cells from various taxonomic groups. By capturing the full emission spectra, spectral cytometry allows precise identification and enumeration of phytoplankton, facilitating a comprehensive understanding of community composition, structure, and dynamics. Additionally, it offers insights into responses to environmental stressors and enhances the characterization of microbiota associated with growing algae, advancing our understanding of phytoplankton ecology, synthetic biology, and evolution.

One of the major challenges in conventional flow cytometry, specifically relevant to phytoplankton research, is the issue of autofluorescence. While in conventional flow cytometry, autofluorescence often interferes with fluorophore emission, and is, therefore, an unwanted signal; spectral cytometry utilizes this phenomenon as a separate parameter(s). In addition to this, the ability to measure the full spectrum of light emissions from each cell allows us to differentiate between various combinations of intrinsic fluorophores that conventional instruments cannot distinguish. This is particularly advantageous in phytoplankton research, due to the high diversity of algal pigments and their overlapping spectral signatures. Alternately, approaches that provide high taxonomic resolution favor methods that allow for genus- to species-level characterization, such as amplicon-sequencing (Sommeria-Klein et al., 2021).

In this work, we demonstrate the effectiveness of full-spectrum cytometry in differentiating between different algal taxa based on their autofluorescent signatures and light scattering parameters. Clear separation between picocyanobacteria (*Synechococcus* sp. and *Synechocystis* sp.), as well as between closely related phytoplankton taxa (Synechococcales order) was demonstrated. Recorded spectral signatures were then utilized in the construction of a spectral library, which is a necessary step in monitoring synthetic communities over time. It was reported before that the spectral shape of the functional absorption spectra is remarkably constrained within major phylogenetic groups of phytoplankton, including diatoms, haptophytes, dinoflagellates, and chlorophytes (Gorbunov et al., 2020) and variability is significantly smaller than the difference between groups. In our study, the emission spectra, obtained from microalgae with a help of ID7000 spectral cell analyzer highlight the ability to discriminate between multiple phytoplankton species in a synthetic mix, even in the presence of closely related taxa. This precision is achieved using optimazed sets of virtual filters and gating strategies, which enable the clear separation of subpopulations. The temporal monitoring of synthetic phytoplankton communities further demonstrates the applicability of full-spectrum cytometry in tracking community dynamics and responses to environmental changes. Comparative analysis with other instruments, such as BD FACSAria cell sorter and ImageStreamX MarkII imaging flow cytometer, demonstrates the higher flexibility of the ID7000 spectral cell analyzer.

Development of biological applications of full spectrum cytometry in phytoplankton research is still in its early stages, with most published work focusing on clinical applications (Wanner et al., 2022; Peixoto et al., 2022; Dott et al., 2024). One of the limitations of this approach relates to the particle size range capabilities of spectral cytometry instrumentation similar to commercially available cytometers. As a result, bulky and lengthy plankton such as many colonial and filamentous cyanobacteria, diatoms, etc. cannot be captured during the analysis. In addition to this, *in situ* measurement of planktonic cells is not available. In contrast, a range of other cytometric instruments, such as FlowCAM and CytoBuoy designed for phytoplankton, have significantly extended particle size range allowing analysis of particles of 500 µm width and over 1000 µm length (Dubelaar et al., 1989; Sieracki et al. 1998; Dubelaar, Gerritzen, 2000). Implementing full-spectrum cytometry in phytoplankton research requires reproducible protocols and standardization throughout the workflow, from sample collection and handling to data acquisition and analysis. Additionally, natural variability in phytoplankton populations, including cell size, shape, and pigment composition differences, requires optimized controls to minimize data variability. With the latest ID7000 software update, system standardization for data reproducibility became available. In the context of phytoplankton research, this means that recorded algal libraries containing unique spectral signatures can be utilized across different experiments, time points and multiple instruments, minimizing data variability. Current work has resulted in a library, which includes spectral signatures of over 30 unique phytoplankton species across different taxa.

The stability of microbial part of artificial algal-microbial mix in accordance with data from other group (Li et al., 2022) reporting population stability with mutualistic co-culturing for up to 10 days. Microorganisms within the consortia could be unculturable, hence culture-independent methods such as high-throughput sequencing and molecular fingerprinting are used for characterization of microbial part of consortia (De Roy et al., 2014). We suggest a nanopore-based sequencing at species-level to be performed not at the end of experiment but in parallel with spectral cytometry evaluation or weekly.

In conclusion, by single-cell analysis and differentiating between different algal taxa based on their autofluorescent spectral signatures and light scattering parameters, full-spectrum cytometry overcomes the limitations of traditional methods such as microscopy, pigment analysis, and conventional cytometry. Its application extends to constructing spectral libraries for various phytoplankton groups and monitoring synthetic communities over time, providing detailed insights into community structure, dynamics, and responses to environmental stressors. The potential for parallel sequencing and associated microbial analysis at the multi-omics level is needed to fully elucidate the microbial functions occurring in artificial algal communities. This technology provides possibilities for a wide range of research applications, from addressing fundamental ecological and evolutionary questions to applied biology, thereby advancing the field of environmental research.

## Authors contributions

A.M. designed the study, performed the experiments, analyzed the data. S.M. and N.F. contributed algal cells, in algal cells characterization. I.A.V. and N.S.B. initiated and supervised the project. A.M. and N.S.B. wrote the manuscript with contributions from all authors.

## Acknowledgements

We would like to acknowledge contribution of Aigul Kussanova during the initial stages of this work and Polina Len during sequence classification. We would also like to thank Dmitry Malashenkov and Aiym Duisen for helpful discussion and input, and Rudolf Bichele and Steffen Boechner from Sony Biotechnology for their instrumental help and support.

## Data availability

Raw sequences obtained during the analysis were deposited in the National Center for Biotechnology Information (NCBI) Sequence Read Archive (SRA) under the BioProject ID PRJNA1131605. Spectral and cytometric data obtained during this study are available from the corresponding author on reasonable request.

