## Supplemental_Data_1 for "Spectral algal fingerprinting and long sequencing in synthetic algal-microbial communities"

**Supplementary Table 1.** List of algal cultures used for recordings during artificial mix experiment and for library construction using ID7000 spectral cell analyzer

| Phylum | Order | Species | Habitat | Strain number |
|---|---|---|---|---|
| Cyanobacteria | Nostocales | *Dolichospermum urugayensis* | Freshwater | 2498 |
| | | *Nodularia sphaerocarpa* | Freshwater | 50.79 |
| | Gloeobacterales | *Gloeobacter violaceus* | Freshwater | 35.87 |
| | Chroococcales | *Microcystis* sp. | Freshwater | CCAC 7146B |
| | Synechococcales | *Synechococcus* sp. | Freshwater | CCAC 2944B |
| | | *Synechocystis* sp. | Freshwater | CCAC 4059B |
| Chlorophyta | Chlorodendrales | *Scherffelia dubia* | Freshwater | CCAC 1398 B |
| | Chlamydomonadales | *Chlamydomonas* sp. | Freshwater | CCAC 4527 B |
| | | *Haematococcus pluvialis* | Freshwater | CCAC 2072B |
| | Chlorellales | *Chlorella vulgaris* | Freshwater | 211-12 |
| | Sphaeropleales | *Acutodesmus obliquus* | Freshwater | 22.81 |
| Cryptophyta | Pyrenomonadales | *Chroomonas* sp. | Freshwater | CCAC 2614B |
| | | *Rhodomonas* sp. | Freshwater | CCAC 1479B |
| | Cryptomonadales | *Cryptomonas* sp. | Freshwater | CCAC 2345B |
| Rhodophyta | Stylonematales | *Chroothece richteriana* | Marine | 104.79 |
| | Porphyridiales | *Porphyridium cruentum cf.* | Marine | CCAC 3771 B |
| Dinophyta | Peridiniales | *Peridinium* sp. | Freshwater | CCAC 1612 B |
| Euglenophyta | Euglenida | *Euglena sanguinea* | Freshwater | CCAC 3518 B |
| Bacillariophyta | Fragilariales | *Fragilaria* sp. | Freshwater | CCAC 5509 B |

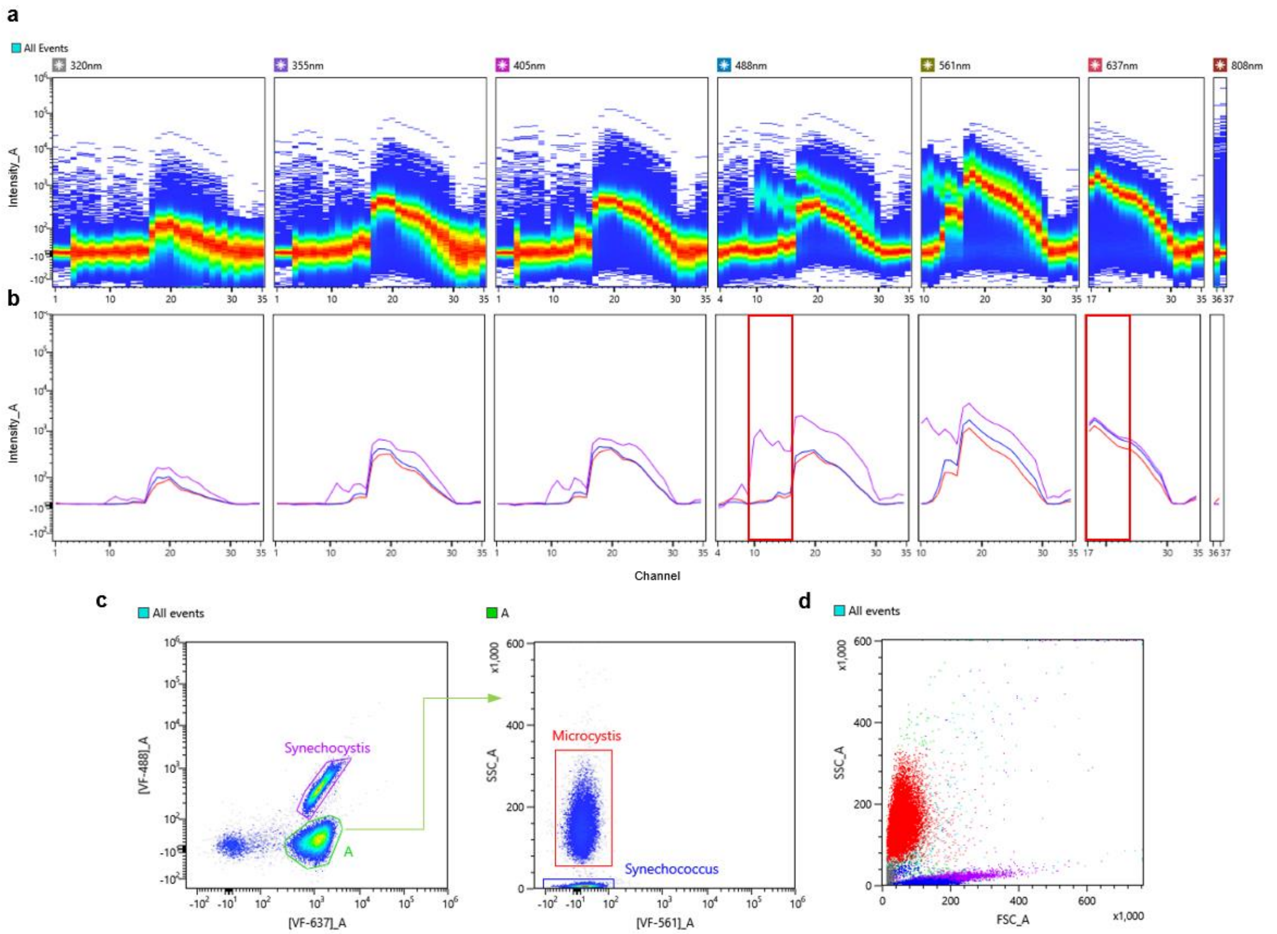


**Supplementary Figure 1.** Discrimination of picocyanobacteria in a synthetic mix based on autofluorescence. **(a)** Ribbon plot demonstrating combined spectral signatures of 3 cyanobacterial representatives across 7 excitation lasers: *Synechococcus* sp., *Synechocystis* sp., and *Microcystis* sp.; **(b)** Overlay plot of individual spectral signatures for each cyanobacterial representative (*Synechococcus* sp. - blue, *Synechocystis* sp. - violet, and *Microcystis* sp. - red); red regions indicate regions of interest, used for setting an optimal pair of VFs; **(c)** Dot plot of **[VF-637]**_A against **[VF-488]_**A with two autofluorescent populations identified: *Synechocystis* sp., population **A** was then plotted with **[VF-561]_**A against **SSC**_A to resolve two subpopulations *of Microcystis* sp. and *Synechococcus* sp.; **(d)** Dot plot demonstrating relative position of each of the identified picocyanobacterial species on FSC_A against SSC_A plot.

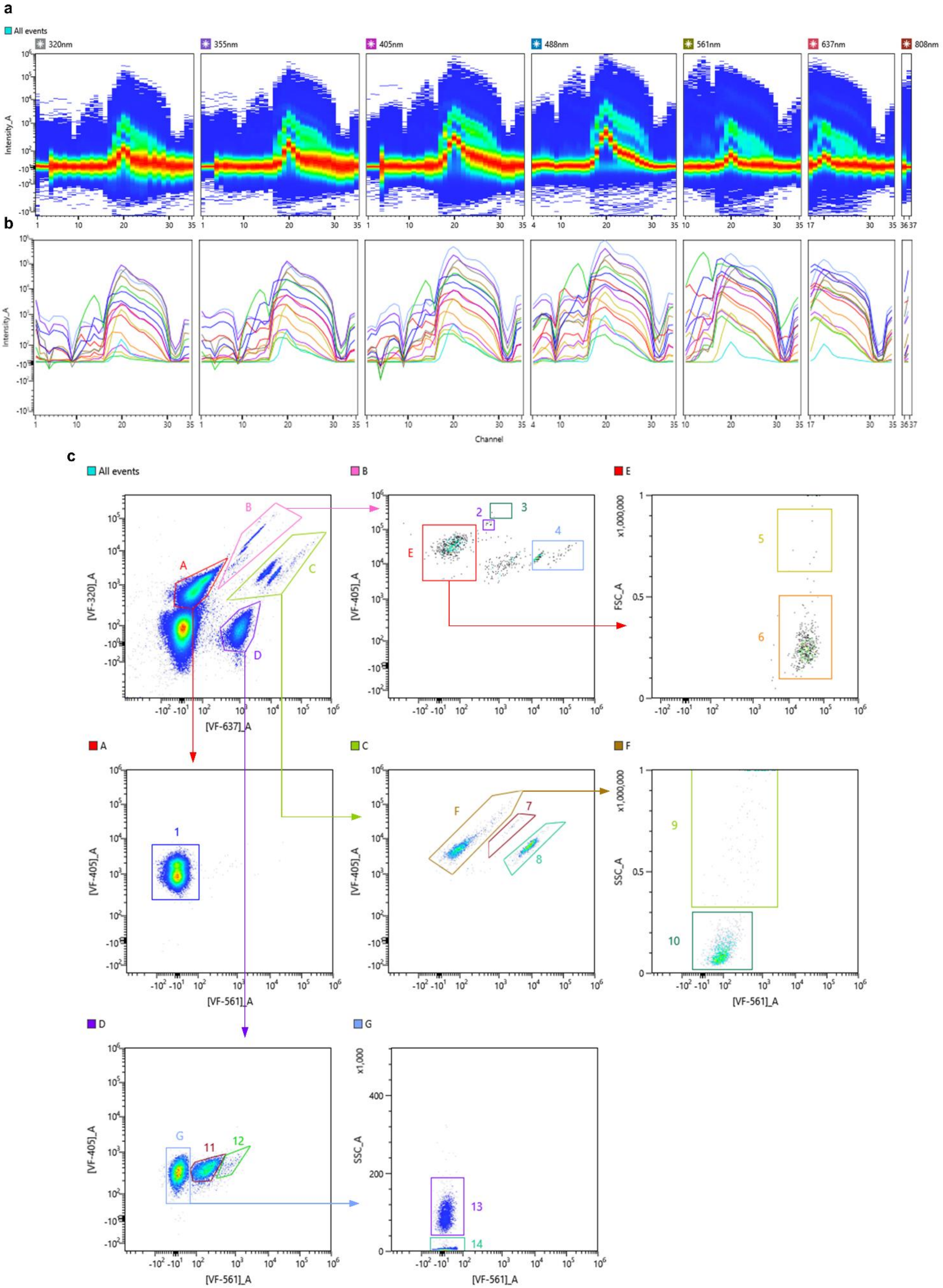
a
All events
320nm
355nm
405nm
488nm
561nm
637nm
808nm
Intensity_A
b
Intensity_A
Channel
c
All events
[VF-320]_A
[VF-637]_A
A
B
C
D
B
[VF-405]_A
E
2
3
4
E
FSC_A
x1,000,000
5
6
A
[VF-405]_A
[VF-561]_A
1
C
[VF-405]_A
F
7
8
F
SSC_A
x1,000,000
[VF-561]_A
9
10
D
[VF-405]_A
[VF-561]_A
G
11
12
G
SSC_A
x1,000
[VF-561]_A
13
14

**Supplementary Figure 2.** Gating strategy for discrimination of 14 groups of phytoplankton species based on autofluorescence in a complex synthetic mix. **(a)** Ribbon plot demonstrating combined spectral signatures of 16 phytoplankton species across 7 excitation lasers. **(b)** Overlay plot of individual spectral signatures for each phytoplankton representative. (**c**) Gating strategy for discrimination of next subpopulation within the synthetic mix: 1 – Chlorophyta (*Chlorella vulgaris* and *Acutodesmus obliquus*), 2 – *Euglena sanguinea*, 3 – *Peridinium* sp., 4 – *Cryptomonas* sp., 5 – *Fragilaria* sp., 6 – *Haematococcus pluvialis*, 7 – *Chroothece richteriana*, 8 – *Porphyridium cruentum* cf., 9 – filamentous Cyanobacteria (*Dolichospermum urugayensis* and *Nodularia sphaerocarpa*), 10 – *Chroomonas* sp., 11 – *Synechocystis* sp., 12 – *Gloeobacter violaceus*, 13 – *Microcystis* sp., 14 – *Synechococcus* sp.

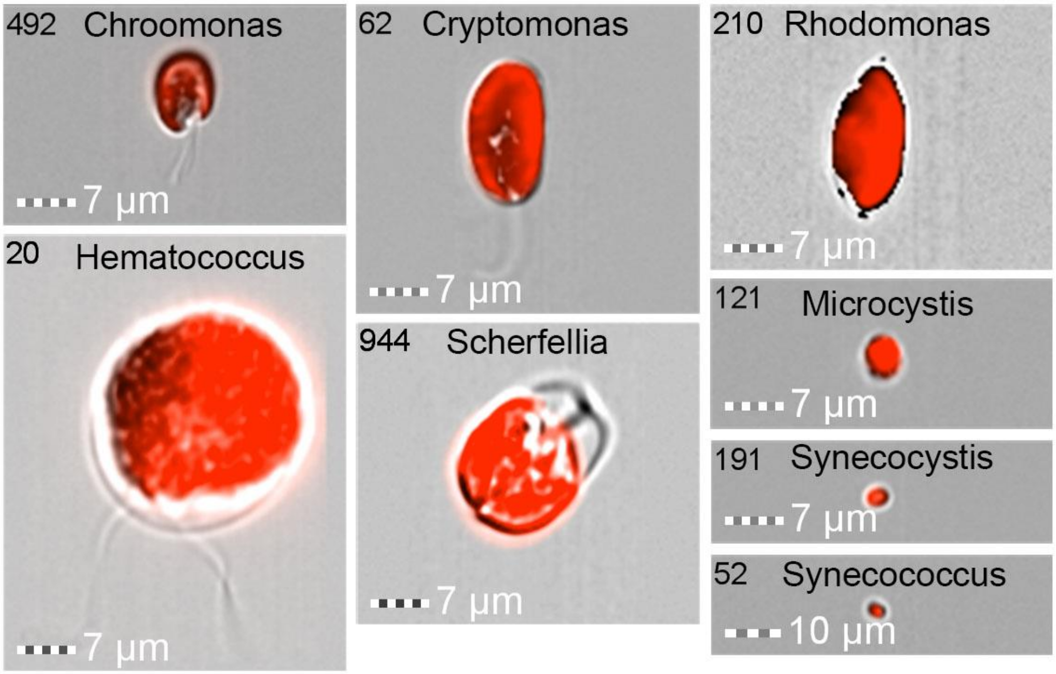


**Supplementary Figure 3.** The gallery of algal images taken with ImageStream MKII (Amnis-Cytek, USA) of phytoplankton species recorded during the artificial mix experiment.

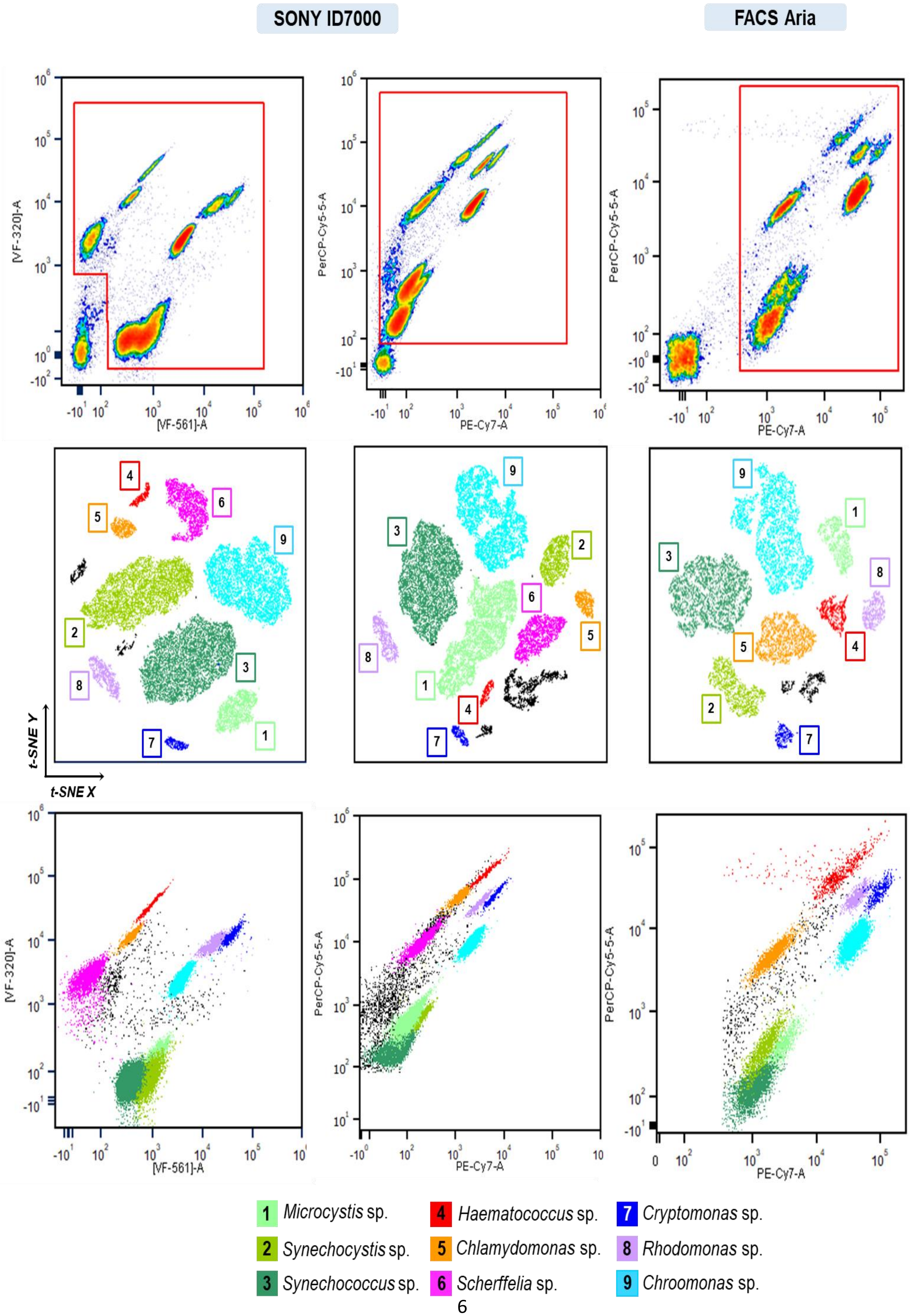
SONY ID7000
FACS Aria
[VF-320]-A
[VF-561]-A
PerCP-Cy5-5-A
PE-Cy7-A
t-SNE Y
t-SNE X
1 Microcystis sp.
2 Synechocystis sp.
3 Synechococcus sp.
4 Haematococcus sp.
5 Chlamydomonas sp.
6 Scherffelia sp.
7 Cryptomonas sp.
8 Rhodomonas sp.
9 Chroomonas sp.

62

**Supplementary Figure 4.** Identification of artificial mix subpopulations compared between ID7000 Spectral Cell Analyzer (SONY Biotechnologies) and BD FACSAria (BD Biosciences). The first column represents data recorded on SONY ID7000 using a set of VFs best optimized for artificial mix discrimination. Opt-SNE demonstrates successful discrimination of the 9 subpopulations within the mix based on VF data. The second column represents data recorded on ID7000 (Sony Biotechnologies) using a set of VFs matching optical filters used on BD FACSAria (BD Biosciences). Similarly, all 9 subpopulations were discriminated. Last column represents data recorded on BD FACSAria. Based on the optical configuration of the instrument, only 8 subpopulations were identified using opt-SNE.

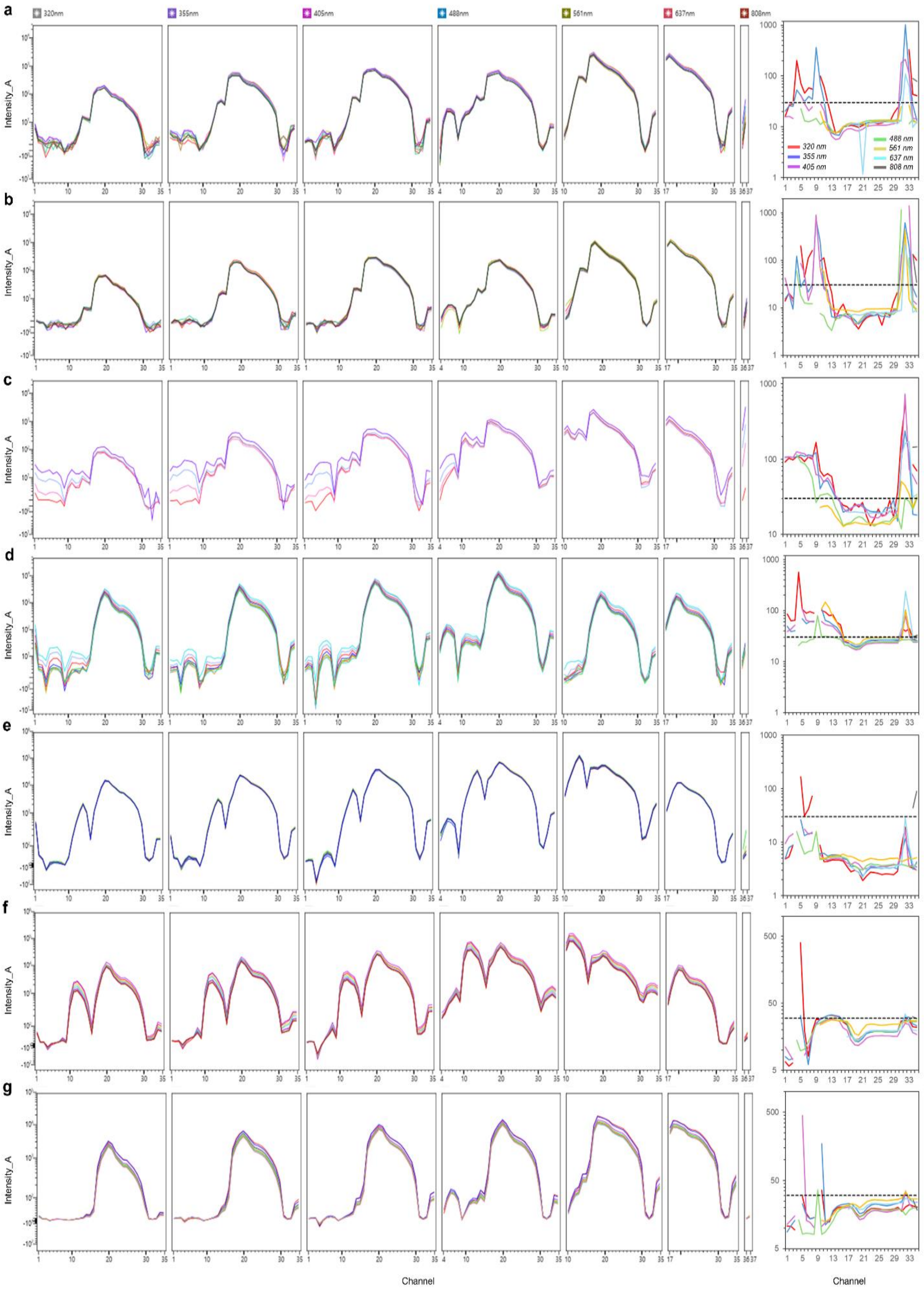
a
b
c
d
e
f
g
320nm
355nm
405nm
488nm
561nm
637nm
808nm
Intensity_A
320 nm
355 nm
405 nm
488 nm
561 nm
637 nm
808 nm
Channel
Channel

73

**Supplementary Figure 5.** Temporal overlay of spectral signatures recorded throughout the experiment. Individual spectral signatures for each member of the synthetic mix (**(a)** – *Microcystis* sp., **(b)** - *Synechococcus* sp., **(c)** – *Synechocystis* sp., **(d)** – *Chlamydomonas* sp., **(e)** - *Cryptomonas* sp., **(f)** – *Rhodomonas* sp., **(g)** – *Chroomonas* sp.) recorded over the period of 9 days overlayed to demonstrate temporal changes in signal intensity. To display signal dispersion, coefficient of variation (CV) was calculated and plotted across all detection channels for each of the excitation lasers. A threshold line at 30% was plotted for additional reference.

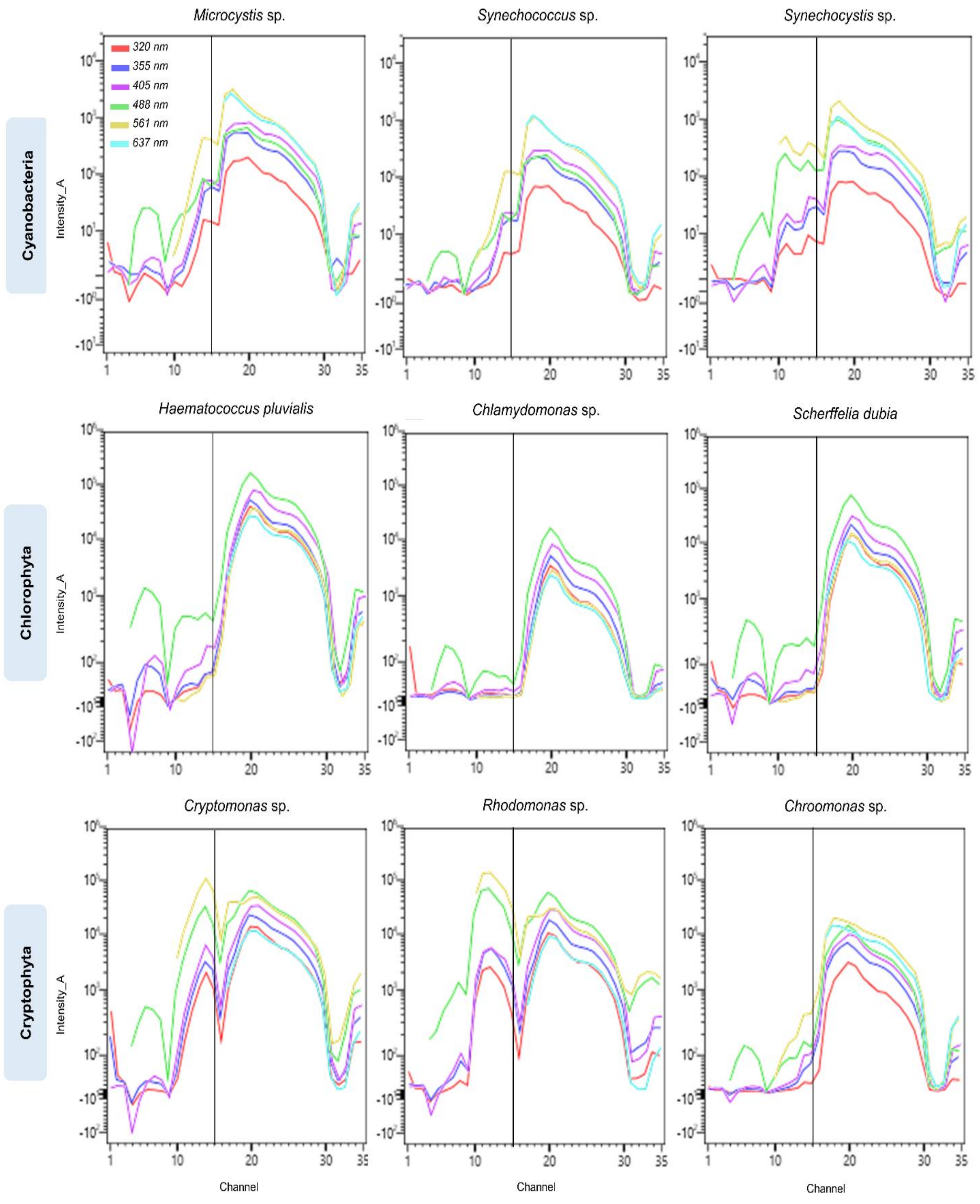


**Supplementary Figure 6.** Overlay plots for each phytoplankton species within a synthetic mix across 7 excitation lasers to demonstrate the variability of emission signal for each excitation source. The first row demonstrates overlay plots for Cyanobacterial representatives, second – Chlorophyta, and third – Cryptophyta. The black line denotes cutoff at channel 15 (624 – 635.3 nm).
